# Error Correcting Optical Mapping Data

**DOI:** 10.1101/285692

**Authors:** Kingshuk Mukherjee, Darshan Washimkar, Martin D. Muggli, Leena Salmela, Christina Boucher

## Abstract

Optical mapping is a unique system that is capable of producing high-resolution, high-throughput genomic map data that gives information about the structure of a genome [21]. Recently it has been used for scaffolding contigs and assembly validation for large-scale sequencing projects, including the maize [32], goat [6], and amborella [4] genomes. However, a major impediment in the use of this data is the variety and quantity of errors in the raw optical mapping data, which are called Rmaps. The challenges associated with using Rmap data are analogous to dealing with insertions and deletions in the alignment of long reads. Moreover, they are arguably harder to tackle since the data is numerical and susceptible to inaccuracy. We develop cOMet to error correct Rmap data, which to the best of our knowledge is the only optical mapping error correction method. Our experimental results demonstrate that cOMet has high prevision and corrects 82.49% of insertion errors and 77.38% of deletion errors in Rmap data generated from the *E. coli* K-12 reference genome. Out of the deletion errors corrected, 98.26% are true errors. Similarly, out of the insertion errors corrected, 82.19% are true errors. It also successfully scales to large genomes, improving the quality of 78% and 99% of the Rmaps in the plum and goat genomes, respectively. Lastly, we show the utility of error correction by demonstrating how it improves the assembly of Rmap data. Error corrected Rmap data results in an assembly that is more contiguous, and covers a larger fraction of the genome.

## Introduction

In 1993 Schwartz et al. developed *optical mapping*, a system for creating an ordered, genome-wide, high-resolution restriction map of a given organism’s genome. Since this initial development, genome-wide optical maps have found numerous applications including discovering structural variations and rearrangements [24], scaffolding and validating contigs for several large sequencing projects [7, 9, 4], and detecting misassembled regions in draft genomes [16]. Thus, optical mapping has assisted in the assembly of a variety of species–including various prokaryote species [18, 29, 30], rice [31], maize [32], mouse [5], goat [7], parrot [9], and *amborella trichopoda* [4]. The raw optical mapping data is generated by a biological experiment in which large DNA molecules cling to the surface of a microscope slide using electrostatic charge and are digested with one or more restriction enzymes. The restriction enzymes cut the DNA molecule at occurrences of the enzyme’s recognition sequence, forming a number of DNA fragments. The fragments formed by digestion are painted with a fluorescent dye, to allow visibility under laser light and a CCD camera. Computer vision algorithms then estimate fragment length from consolidated intensity of fluorescent dye and apparent distance between fragment ends.

The resulting data from an experiment are in the form of an ordered series of fragment lengths [33]. The data for each single molecule produced by the system is called an *Rmap*. Rmap data has a number of errors due to the experimental conditions and system limitations. In an optical mapping experiment, it is unlikely to achieve perfectly uniform fluorescent staining.

This leads to an erroneous estimation of fragment sizes. Also, restriction enzymes often fail to digest all occurrences of their recognition sequence across the DNA molecule. This manifests as missing restriction sites. Additionally, due to the fragile nature of DNA, additional breaks can incorrectly appear as restriction sites. Lastly, the limitations of the imaging component of the optical mapping system and the propensity for the DNA to ball up at the ends introduces more sizing error for smaller fragments. Interested readers will find more details about the causes of these errors in Valouev *et al*. [26] and Li *et al*. [12]. Because of all these experimental conditions, Rmap data generated through optical mapping experiment has insertion (added cut sites) and deletion (missed cut sites) errors along with fragment sizing errors.

In most applications of optical map data, the Rmaps need to be assembled into a genome wide optical map. This is because the single molecule maps need redundant sampling to overcome the presence of the aforementioned errors, and because single molecule maps only span on the order of 500 Kbp [26]. The first step of this assembly process involves finding pairwise alignments amongst the Rmaps. In order to accomplish this, the challenge of dealing with missing fragment sizes has to be overcome. This challenge is analogous to dealing with insertions and deletions in the alignment of long reads [2]— in fact, it is arguably harder since the data is numerical. At present, the only non-proprietary algorithmic method for pairwise alignment of Rmaps is the dynamic programming based method of Valouev *et al*. [26] which runs in *O*(α*×*β) time where α and β are the number of fragments in the two Rmaps being aligned. To align an optical map dataset containing *n* Rmaps, the complexity becomes *O*(*n*^2^ *×* 𝓁^2^) where 𝓁 is the average size of an Rmap.

This method is inherently computationally intensive but if the error rate of the data could be improved then non-dynamic-programming based methods that are orders of magnitude faster such as Twin [15], OMBlast [10], and Maligner [13] could be used for alignment. This would greatly improve the time required to assemble Rmap data. Thus, we present cOMet in order to address this need. To the best of our knowledge, it is the first Rmap error correction method. Our experimental results demonstrate that cOMet has high precision and corrects 82.49% of insertion errors and 77.38% of deletion errors in Rmap data generated from the *E*.*coli* K-12 reference genome. Out of the deletion errors corrected, 98.26% are true errors. Similarly, out of the insertion errors corrected, 82.19% are true errors. Furthermore we show that the assembly of Rmaps is more contiguous and covers a larger fraction of the genome if the Rmaps are first error corrected. It also successfully scales to large genomes, improving the quality of 78% and 99% of the Rmaps in the plum and goat genome, respectively.

## Background

From a computer science perspective, optical mapping can be seen as a process that takes in two strings: a nucleotide sequence *S*_*i*_[1, *n*] and a restriction sequence *B*[1, *b*], and produces an array (string) of integers *R*_*i*_[1, *m*]. The array *R*_*i*_ is an Rmap corresponding to *S*_*i*_ and contains the string-lengths between cuts produced by *B* on *S*_*i*_. Formally, *R*_*i*_ is defined as follows: *R*_*i*_[*j*] = *y* – *x* where *y* represents the location(starting index) of *j*^*th*^ occurrence of *B* in *S*_*i*_ and *x* represents the location of (*j* – 1)^*th*^ occurrence of *B* in *S*_*i*_ and *R*_*i*_[1] = *y* – 1 and *R*_*i*_[*m*] = *n* – *x*. For example, say we have *B* = *act* and *S*_*i*_ = *atacttactggactactaaact*. The locations of *B* in *S*_*i*_ are as follows: 3,7,12,15,20. Then *R*_*i*_ will be represented as *R*_*i*_ = 2, 4, 5, 3, 5, 2. The size of an Rmap denotes the number of fragments in that Rmap. Therefore the size of *R*_*i*_ is 6.

We note that millions of Rmaps are produced for a single genome since optical mapping is performed on many cells of the organism and each cell provides thousands of Rmaps. The Rmaps can be assembled to produce a genome wide optical map. This is analogous to next generation shotgun sequencing where Rmaps are analogous to reads and a genome-wide optical map is analogous to the assembled whole genome.

There are three types of errors that can occur in optical mapping: (1) missing cut sites which are caused by an enzyme not cleaving at a specific site, (2) additional cut sites which can occur due to random DNA breakage and (3) inaccuracy in the fragment size due to the inability of the system to accurately estimate the fragment size. Continuing again with the example above, a more representative example Rmap would include these errors, such as 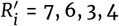.

The error rates of optical maps depends on the platform used for generating the maps. A recent paper by Li *et al*. [12] studied the error rates of optical maps produced by the Irys system from BioNano Genomics. According to their study, a missing cut site type of error i.e., error type (1) happens when a restriction site is incompletely digested by the enzyme and causes two flanking fragments to merge into one large fragment. The probability of complete digestion of a restriction site can be modeled as a Bernoulli trial whose probability of success is a function of the size of the two flanking fragments. Additional cut sites i.e., error type (2) results from random breaks of the DNA molecule. The number of false cuts per unit length of DNA follows a Poisson distribution. The inaccuracy of the fragment sizes, i.e., error type (3), is modeled using a Laplace distribution. If the observed and actual size of a fragment are *o*_*k*_ and *r*_*k*_ respectively, then the sizing error is defined as *s*_*k*_ = *o*_*k*_/*r*_*k*_ and

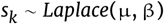

where µ and β -the parameters of the laplace distribution -are functions of *r*_*k*_. In practice, when aligning a pair of Rmaps, one should allow for twice the error rate of a single Rmap since each Rmap will deviate from the genomic map by the above parameters.

Valouev *et al*. [26] provides a dynamic programming algorithm for pairwise alignment, which generates a score for every possible alignment between two Rmaps and returns the alignment which achieves the highest score, which is referred to as the *S*-score. It is computed within a standard dynamic programming framework, similar to Smith-Waterman alignment [23]. The scoring function is based on a probabilistic model built on the following assumptions: the fragment sizes follow an exponential distribution, the restriction sites follow an independent Bernoulli process, the number of false cuts in a given genomic length is a Poisson process, and the sizing error follows a normal distribution with mean zero and variance following a linear function of the true size. Lastly, a different sizing error function is used for fragments less than 4 kbp in length since they do not converge to the defined normal distribution. The score of an alignment is calculated as the sum of two functions; one function that estimates and scores the sizing error, and a second that predicts and scores the presence of additional and/or missing cut sites between the fragments. The *S*-score will be used later in this paper to evaluate the error correction process.

## Methods

Given a set of *n* Rmaps R = {*R*1, .., *R*_*n*_} our method aims to detect and correct all errors in R by considering each *R*_*i*_ *∈* R and finding a set of Rmaps that originate from the same part of the genome as *R*_*i*_. This step is performed heuristically in order to avoid aligning every pair of Rmaps in R.

### Preprocessing

Our first step is to remove the first and last fragments from each Rmap in R. These fragments have one of their edges sheared by artifacts of the DNA prep process (preceding the optical mapping process) and not by restriction enzymes. Unless removed, they can misguide alignment between two Rmaps during the error correction process. In addition, short Rmaps, i.e., those that have less than 10 fragments, are removed at this stage since any Rmap that contains less than 10 fragments is typically deemed too small for analysis even in consensus maps [1]. Next, the data is quantized so that a given genomic fragment is represented by the same value across multiple Rmaps despite the noise. Our quantization method assigns a unique value to a range of fragment sizes by dividing each fragment size by a fixed integer, denoted as *b*, and rounding to the nearest integer. For example, if an Rmap *R*_*i*_ = {36, 13, 15, 20, 16, 5, 21, 17} is quantized using *b* = 3 then the quantized Rmap will be 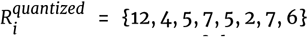.Say another Rmap, *R*_*j*_ = {17, 23, 34, 12, 14, 21, 14, 5} has overlap with *R*_*i*_; however, due to noise in the data, this relation is not apparent. By quantizing *R*_*j*_ using the same *b* = 3 we get 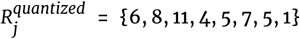. This allows us to uncover a region (in this case {4, 5, 7, 5}) which is common to both the Rmaps. It should be noted that in some cases, a fragment may have different values across two Rmaps even after quantization (for example the fragment values 36 from *R*_*i*_ and 34 from *R*_*j*_ are quantized to 12 and 11 respectively). The quantized data is used to find the set of *related* Rmaps as explained in the next section.

The setting of parameter *b* depends on the amount of sizing error in the optical map data. With zero sizing error *b* can be set at 1. As sizing error increases, the value of *b* is increased accordingly. If the value of *b* is too small, we are not be able to uncover relations between overlapping Rmaps and if the value is too large then unrelated Rmaps have common regions in their quantized states — which makes them appear related. Considering the error rate of optical maps from BioNano genomics, the default value of *b* = 4000.

### Finding Related Rmaps

We refer to two Rmaps as *related* if their corresponding error-free Rmaps originate from overlapping regions of the genome. Next, we define a *k*-mer as a string of *k* consecutive fragments from a (quantized) Rmap. For example if we have the Rmap *R* = {3, 3, 5, 2, 6, 5, 5, 1} and *k*=4 then the following *k*-mers can be extracted from *R*: (3,3,5,2), (3,5,2,6), (5,2,6,5), (2,6,5,5) and (6,5,5,1). In order to avoid aligning all pairs of Rmaps to find the related Rmaps, we use the number of common *k*-mers to discriminate between pairs of Rmaps that are related and those that are not. To accomplish this eficiently, we first extract all unique *k*-mers in each quantized Rmap, and construct a hash table storing each unique *k*-mer as a key and the list of Rmaps containing an occurrence of that *k*-mer as the value. We call this the *k*-mer index. Next, we consider each *R*_*i*_ in R and use the *k*-mer index to identify the set of Rmaps that have *m* or more *k*-mers in common with *R*_*i*_. Unfortunately, this set, although it contains all related Rmaps, it also likely contains Rmaps that are not related to *R*_*i*_. Therefore, we filter this set of Rmaps using a simple heuristic that tries to match each Rmap in this set with *R*_*i*_ in order to ascertain if it is related to *R*_*i*_. The heuristic traverses through two Rmaps (*R*_*i*_ and one Rmap from the set, say *R*_*j*_) attempting to match subsets of the fragments from each until it either reaches the end of one Rmap or it fails to match the fragments. We start the traversal from the first matching *k*-mer between *R*_*i*_ and *R*_*j*_. We denote the position of the next fragment to be matched in *R*_*i*_ and *R*_*j*_ as *x* and *y*, respectively, and assume that each fragment prior to these positions is matched. Next, we consider all combinations of matching the fragments at positions *x, x* + 1 and *x* + 2 of *R*_*i*_ with fragments at positions *y, y* + 1 and *y* + 2 of *R*_*j*_. We evaluate the cost of each combination based on the difference in the total size of fragments from *R*_*i*_ and *R*_*j*_. That is *∀*α, β = [0, 2],

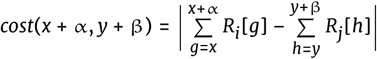

where *R*_*i*_[*g*] and *R*_*j*_[*h*] denotes the *g*-th and *h*-th fragments of *R*_*i*_ and *R*_*j*_ respectively. We select the combination with the least cost; if there exists a tie, we select the match that has the least number of added or missing cut sites (That is, the combination with the least value of α + β). If this selected match leads to a cost that is greater than a specified threshold (which was set to 25% of the larger sized fragment in practice), then we conclude that there is not a match at these positions and return that *R*_*i*_ and *R*_*j*_ are unrelated. Otherwise, we increment *x* and *y* accordingly and move onto the next fragments. If this heuristic continues until the last fragment of either *R*_*i*_ or *R*_*j*_ is reached then we return that *R*_*i*_ and *R*_*j*_ are related. Using this heuristic we filter out the Rmaps that were deemed to be related based on the number of *k*-mers in common with *R*_*i*_ but are infact unrelated to *R*_*i*_.

The setting of parameters *k* and *m* are correlated. If the value of *k* is increased, that makes the *k*-mers more specific, hence, the value of *m* is lowered. On the other hand, if the value of *k* is reduced, then we increase the value of *m*. The value of *k* is increased when there are fewer insertion and deletion errors and decreased otherwise. The default values are *k* = 4 and *m* = 1.

### Rmap Alignment

Next, for each *R*_*i*_ in R, we use the alignment method of Valouev *et al*. [26] *to find the S*-score of all pairwise alignments between *R*_*i*_ and each Rmap in its set of related rmaps. The Rmaps that have an alignment score, i.e., *S*-score less than a defined threshold (which we denote as *S*_*t*_), are removed from the set of related Rmaps and the alignments of the remaining Rmaps are stored in a *multiple alignment grid*, denoted as A_*i*_. This grid is a two-dimensional array of integer pairs, where the number of rows is equal to the number of remaining Rmaps in the set of related Rmaps of *R*_*i*_ and the number of columns is equal to the number of fragments in *R*_*i*_. An element of this array, A_*i*_[*j, k*] stores an integer pair in the form of (*x, y*) representing that *x* fragments of *R*_*i*_, (which includes the *k*-th fragment of *R*_*i*_) matches to *y* fragments of *R*_*j*_ in the optimal alignment between *R*_*i*_ and *R*_*j*_. Figure 1 illustrates an example of A_*i*_. The first fragment of *R*_*i*_ does not match with any fragment of *R*_*j*_ and therefore, (0, 0) is stored at this position. Fragments 2, 5, 6, 8 and 9 of *R*_*i*_ each matches with one fragment of *R*_*j*_, e.g., 1, 3, 4, 7 and 8, respectively. To represent these matches, we store a (1,1) in 2nd, 5th, 6th, 8th and 9th column of row *j*. Fragments 3 and 4 of *R*_*i*_ match with one fragment of *R*_*j*_, i.e., the 2nd fragment. To represent this, we store (2,1) in A_*i*_[*j*, 3] and A_*i*_[*j*, 4]. Fragment 7 of *R*_*i*_ matches with two fragments of *R*_*j*_, i.e., the 5th and 6th fragments. To represent this, we store (1,2) in A_*i*_[*j*, 7]. Fragments 10 and 11 of *R*_*i*_ match with two fragments of *R*_*j*_, i.e., the 9th and 10th fragments. To represent this, we store (2,2) in positions A_*i*_[*j*, 10] and A_*i*_[*j*, 11]. Finally, fragments 12 and 13 match with three fragments of *R*_*j*_, i.e., fragments 11, 12 and 13. In this case, we store (2,3) in positions A_*i*_[*j*, 12] and A_*i*_[*j*, 13].

**Figure 1.**
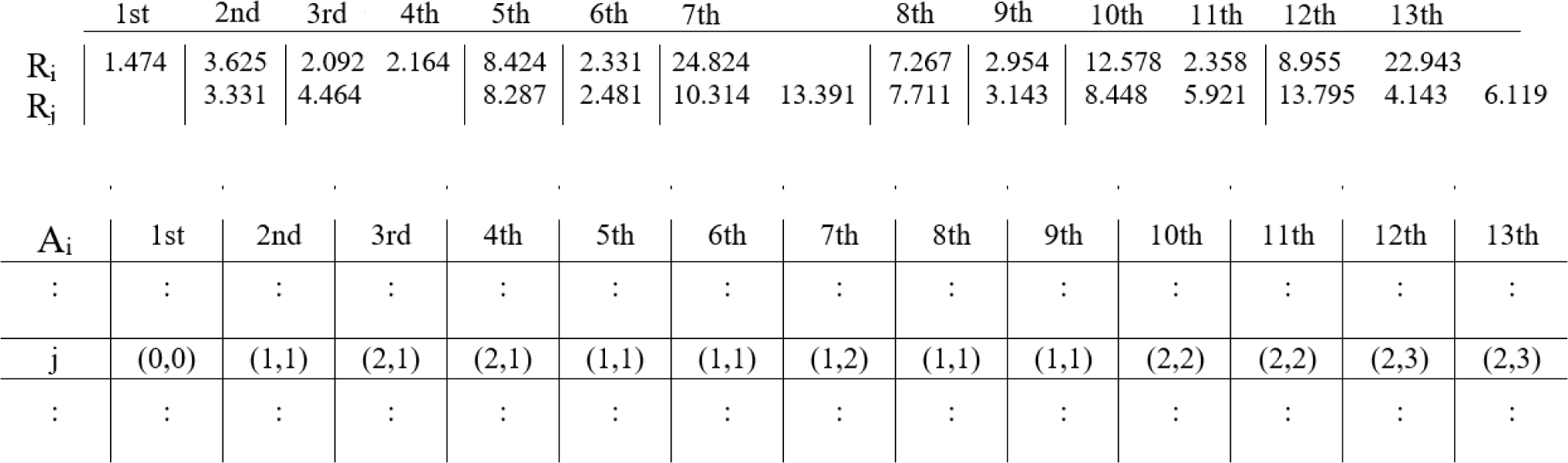
An alignment between *Ri* and *Rj* as given by Valouev *et al*. [26] and its corresponding entry in the multiple alignment grid A*i. Each column of* A*i* represents one fragment from *Ri* and each row represents one Rmap from its’ set of related Rmaps. The fragment sizes are in Kbp.

The setting of parameter *S*_*t*_ controls the number of Rmaps that are included in the multiple-alignment-grid of an Rmap. If we increase the value of *S*_*t*_, fewer Rmaps will be added to the grid — but the ones included will be of higher quality (i.e. have greater overlap with the Rmap under consideration). The default value for the parameter *S*_*t*_ = 8. We show in the experiment section how we select this value.

### Error Correcting Using the Consensus

The multiple alignment grid is used to find the consensus grid, denoted as C_*i*_, for Rmap *R*_*i*_. The grid C_*i*_ is a one-dimensional array of integer pairs with size equal to the number of fragments in *R*_*i*_. The grid is constructed for each *R*_*i*_ in R by iterating through each column of A_*i*_ and finding the most frequent integer pair, breaking ties arbitrarily. The most frequent integerpair is stored at each position of C_*i*_ if the frequency is above a given threshold *d*; otherwise, (0,0) is stored. Figure 2 illustrates the construction of a consensus grid from an alignment grid. The type of error in each fragment of *R*_*i*_ can be identified using C_*i*_[*k*] = (*x, y*) as follows: if *x* and *y* are equal then a sizing error occurs at the *k*-th fragment of *R*_*i*_, otherwise, if *x* is greater than *y* then an additional cut site exists, and lastly, if *x* is less than *y* then a missing cut site exists. Next, we use C_*i*_ and A_*i*_ to correct these errors in *R*_*i*_. For each fragment of *R*_*i*_, we consider the consensus stored at the corresponding position of C_*i*_, identify the positions in the corresponding column of A_*i*_ that are equal to it, and replace the fragment of *R*_*i*_ with the mean total fragment size computed using the values at those positions in A_*i*_. If C_*i*_ is equal to (0,0) at any position then the fragment at that position in *R*_*i*_ remains unchanged since it implies that there is no definitive result about the type of error in that position. In addition, if consecutive positions in C_*i*_ are discordant then the fragments in those positions in *R*_*i*_ also remains unchanged. For example, if there is a (2,1) consensus at some position of C_*i*_, then we expect the preceding or successive position to also have a (2,1) consensus. However, if this is not the case, then we do not error correct those fragments since the consensus is discordant at those positions. Figure 2 shows this error correction. As it is illustrated, to error correct the second fragment of *R*_*i*_, we compute the average of the matched fragments from related Rmaps 2, 3, 4, 5 and 6 and replace the second fragment of *R*_*i*_ with that value as shown in Figure 2. Similarly, to correct the third fragment in this example, we identify that (2,1) is in the consensus, which implies that majority of the related Rmaps are such that two fragments of *R*_*i*_ match with one fragment from the set of related Rmaps, and therefore, replace the third and fourth fragments with the average from the corresponding Rmaps and positions.

**Figure 2.**
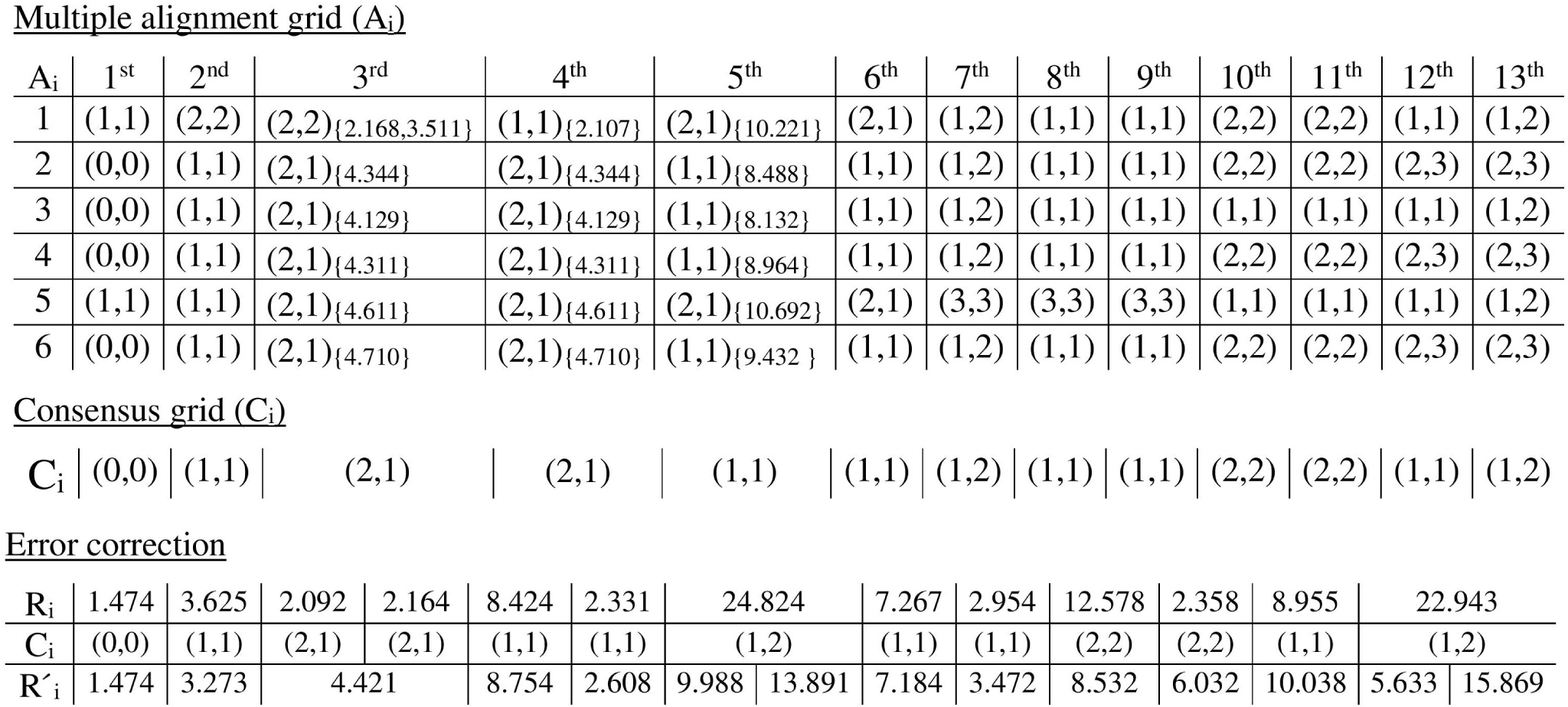
Example of multiple alignment grid and consensus grid. The figure shows the multiple alignment grid A*i* for an Rmap *Ri* and its consensus grid C*i*. Each row of the multiple alignment grid represents the alignment of *Ri* with one of its related Rmaps while the columns represents the fragments of *Ri*. The figure also demonstrates error correction using the consensus grid, with the error corrected Rmap denoted as 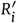. The fragment sizes are in Kbp. To demonstrate the error correction process for the 3rd, 4th and 5th fragments, we also include the fragments (in parentheses) to which they align. The error corrected fragment is the mean of the fragments from the corresponding positions which have the same alignment as the consensus. For example for the 5th fragment, the consensus is (1,1). Therefore the mean of the aligned fragments with (1,1) alignment i.e. 8.488, 8.132, 8.964 and 9.432 is the error-corrected value for the 5th fragment.

The threshold *d* determines the accuracy and precision of error correction. A high value of *d* improves precision but lowers accuracy as many fragments are left uncorrected. Similarly, low value of *d* improves accuracy but lowers precision. The default setting is *d* = 3.

### Complexity

We define 𝓁 to be the length of the longest Rmap in R. Quantization of the Rmaps takes *O*(𝓁 *× n*) time. Constructing the *k*-mer index also takes *O*(𝓁 *× n*) time. The *k*-mer index stores the occurances of each quantized *k*-mer across all Rmaps. Let *u* be maximum frequency of a *k*-mer. That is, a *k*-mer occurs in max *u* Rmaps (in practice *u* << *n*). Then the complexity of finding related Rmaps from the *k*-mer index is *O*(*n×* 𝓁 *× u*). For each Rmap, the filtering heuristic runs in time linear to the size of the Rmap. Therefore, filtering the set of related Rmaps also takes linear *O*(𝓁 *× n*) time. The most expensive step is the pair-wise alignment which uses the Valouev aligner. As mentioned earlier, this aligner is based on DP and therefore has a *O*(𝓁^2^) time complexity to perform one pairwise alignment. If the maximum cardinality of the set of related Rmaps for any Rmap is *v*, then the total complexity of this step is bounded by *O*(*n × v ×* 𝓁^2^). The value of *v* depends on the coverage of the optical map data. The alignment generated using Valouev *et al*. method is stored in the multiple alignment grid in constant time and it takes *O*(*n × v ×* 𝓁) time to generate the consensus maps for *n* Rmaps and error correct them. Thus, the runtime of cOMet is *O*(*n × v ×* 𝓁^2^).

### Datasets

We perform experiments on both simulated and real data. For the real data, we used the Rmap data from the plum [28] and domestic goat [7] sequencing projects. These datasets were built on the OpGen mapping platform and are more error-prone. We also experimented on a human dataset [22] built on the new BioNano platform. This dataset is built using the latest optical mapping technology and has significantly better quality than the plum and goat genomes. The genome size and number of Rmaps for these species are shown in Tables 1 and 1.

**Table 1.** Summary of the real and simulated data. Rmaps with less than 10 fragments were omitted from all the experiments. cOMet was ran on the remaining 2,504, 548,779 and 3,049,439 Rmaps for the E. coli, plum and goat genomes, respectively..

| Genome | Size | No. of Rmaps |
| --- | --- | --- |
| *E. coli* | 4.6 Mbp | 2,504 |
| Plum | 284 Mbp | 749,895 |
| Goat | 2.66 Gbp | 3,447,997 |

**Table 2.** Summary of the real and simulated BioNano data. OM-Sim was used to simulate eight different BioNano datasets; each of which had varying error rates and thus, had a different number of Rmaps.

| Genome | Size | No. of Rmaps |
| --- | --- | --- |
| *E. coli* | 4.6 Mbp | 123,251 - 157,743 |
| Human | 3.2 Gbp | 793,199 |

In addition, we simulated Rmap data from *E. coli* K-12 sub-str. MG 1655 as follows: first, the reference genome was copied 200 times and then uniformly distributed random loci were selected for each of these copies. These loci form the ends of single molecule that would undergo *in silico* digestion. Next, molecules smaller than 150 Kbp were discarded and the cleavage sites for the RsrII enzyme were then identified within each of these simulated molecules. This error free Rmap data is used for validating the output of our method. Lastly, deletion, insertion and sizing errors were incorporated into the error-free Rmaps according to the error model discussed in Li *et al*. [12]. The error model was described earlier in the Background section. This simulation resulted in 2,505 Rmaps, containing 7,485 deletion and 554 insertion errors.

Lastly, we simulated optical map data from a simulation software called OMSim [14] that generates synthetic optical maps which mimics real Bionano Genomics data. The software takes two parameters as input: the False Positive Rate (FP), which is the number of additional cut sites erroneously inserted per 100kbp, and the False Negative Rate (FN), which is the percentage of times a cut site is missed. Using this method, we simulated eight datasets of Rmaps from *E. coli* K-12 substr. MG 1655 using the restriction enzyme BspQI. The default FP and FN rates for BspQI are 1 and 15% respectively.

We generated additional datasets with the following error rates (FP,FN) : (0.5,15%), (1.0,15%), (2.0,15%),(5.0,15%), (1.0,5%), (1.0,25%),(2.0,5%) and (2.0,25%).

### Experiments and Discussion

We performed all experiments on Intel E5-2698v3 processors with 192 GB of RAM running 64-bit Linux. The input parameters to cOMet include: *b* (quantization bucket size), *k* (*k*-mer value), *m* (the number of *k*-mers needed to be conserved between two Rmaps) and *d* (the minimum number of Rmaps required to form consensus at a position). The default parameters are *b*=4000, *k*=4, *m*=1 and *d*=3, and led to the best result across all datasets.

### Determining the value of *S*_*t*_

The setting of the parameter *S*_*t*_ depends on the sensitivity of the Valouev aligner. If the alignment score between two Rmaps is less than *S*_*t*_, then the aligned Rmaps are deemed to be unrelated. We say an Rmap, *R*_*s*_ is *overlapping* with an Rmap, *R*_*t*_ if at least 50% of *R*_*s*_ overlaps with *R*_*t*_. That is, either the first half or the second half of *R*_*s*_ is entirely and exactly (exact fragment matches) contained in *R*_*t*_

We carried out the following experiment to determine the optimum setting for *S*_*t*_. From the set of simulated error-free Rmaps, we computed the set of overlapping Rmaps for each Rmap. We denote this set as *related Rmaps*. Then we used the Valouev aligner to score all pairwise wise alignments between the simulated Rmaps (with errors added) and plot the scores in form of a histogram which is shown in Figure 3. The percentage of related Rmaps with S-score less than 8 is 6.06%. Hence we choose the setting of *S*_*t*_ = 8.

**Figure 3.**
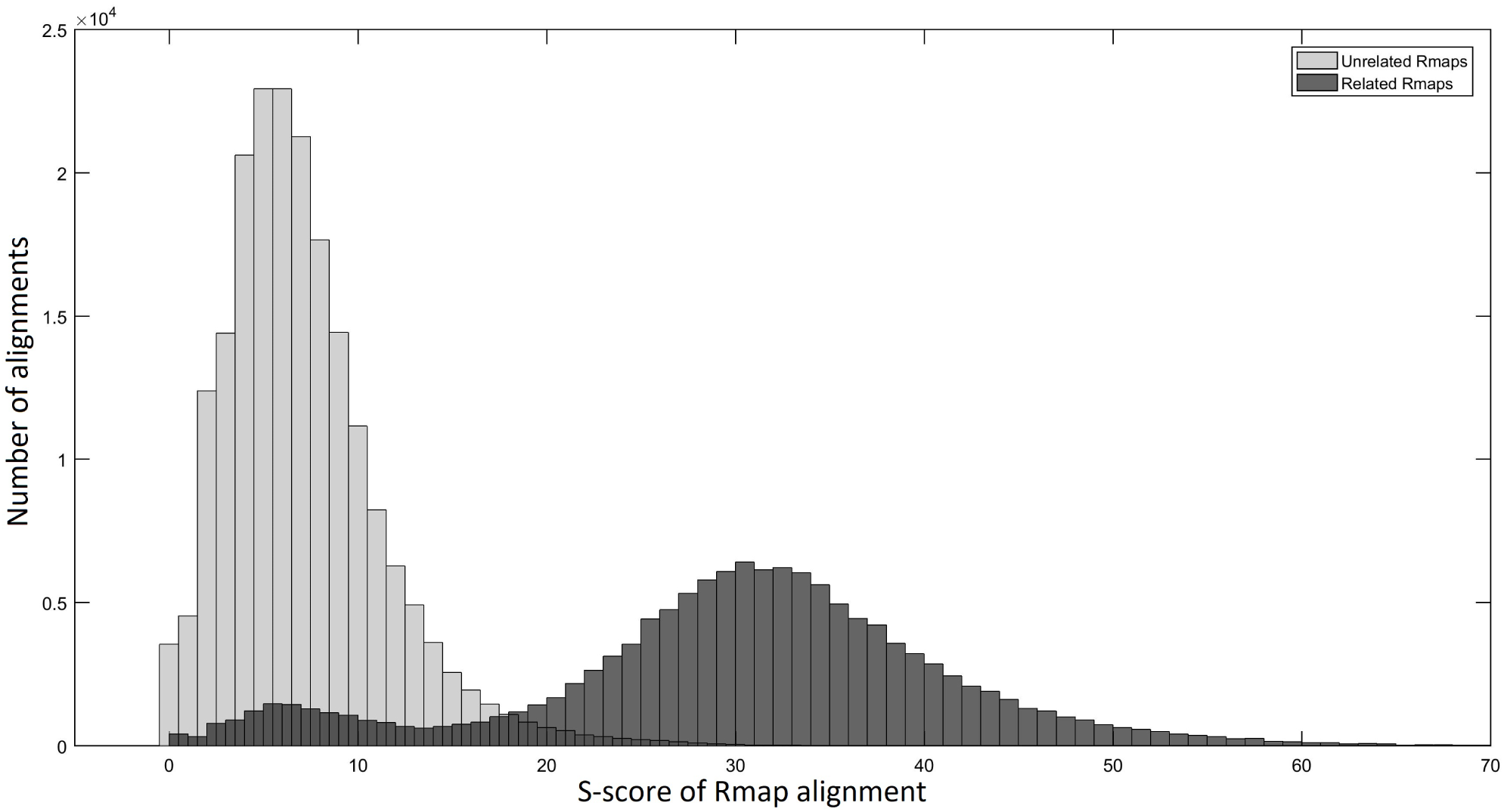
Distribution of S-scores of Rmap alignments between related Rmaps and unrelated Rmaps. The percentage of Related Rmaps with S-score less than 8 is 6.06%. Therefore we choose *St* = 8.

### Experiments with our Simulated Data

The cOMet error correction was ran on the simulated *E. coli* data. The corrected Rmaps were then aligned to the error-free Rmaps to determine the number of corrected insertions and deletions. The results of this experiment are shown in Table 4. To determine the quality of error correction, we computed the true positive rate (TPR), which is the ratio between the number of insertion (or deletion) errors that cOMet correctly identified and removed and the number of insertion (deletion) errors, and the false positive rate (FPR), which is the ratio between the number of insertion (or deletion) errors that cOMet incorrectly identified and removed, and the total number of fragments not containing an insertion (deletion) error. The TPR is 82.49% and 77.38% with respect to the number of corrected insertions and deletion errors; whereas, the FPR is 0.21% and 0.25% with respect to the number of corrected insertions and deletion errors. This demonstrates the high accuracy of the correction made by cOMet. Our method also has high precision. Out of the deletion errors corrected, 98.26% are true errors. Similarly, out of the insertion errors corrected, 82.19% are true errors.

Additionally, for each corrected Rmap we computed the alignment *S*-score of both the original Rmap and the corrected Rmap with the error-free Rmap. We found that for 96.5% of the Rmaps, the *S*-scores improved after error correction. In other words, cOMet brought 96.5% Rmaps closer to their error-free state. The mean *S*-score before error-correction was 44.91 and it improved by 14.03% to 51.30 after error correction. For 17.5% of the Rmaps, (415 Rmaps) the *S*-score improved by more than ten. Lastly, we mention that the error correction was achieved in 241 CPU seconds and using 79.54 MB of memory.

To demonstrate the importance of error correction, we assembled the Rmaps before and after error correction using the Valouev assembler [25]. Table 3 summarizes the results of this experiment. We assembled the uncorrected data into five assembled optical maps and the error-corrected data into two assembled optical maps. The N50 statistic of the assembly increased from 1,242 Kbp for the uncorrected data to 3,348 Kbp for the corrected data. Next, we aligned each assembled map to the genome-wide (error-free) optical map using the Valouev aligner in order to locate their positions on the genome and calculate the percentage of the genome that was covered by at least one of the assembled maps. The genome fraction covered by the five assembled maps from the uncorrected Rmaps was 80%; while the genome fraction covered by the two assembled maps from the corrected Rmaps was 82%. Moreover, the assembled maps from the uncorrected data had 47 insertion and deletion errors when aligned to the reference while the error corrected data had only 34 such errors. In order to further contextualize these results, we assemble the error-free Rmap dataset and summarize this assembly in Table 3.

**Table 3.** Assembly results of Uncorrected Rmaps, Corrected Rmaps and Error-free Rmaps using the Valouev assembler. The Rmaps are simulated from the *E. coli* genome. Each assembled map is aligned to the reference genome-wide (error-free) optical map using the Valouev aligner. The genome-wide optical map contains 383 fragments.

| Rmap status | Assembler Map_id | Number of fragments | Map length (in Kbp) | Alignment location in reference (start-loci, end-loci) |
| --- | --- | --- | --- | --- |
| Uncorrected Rmaps | Assembled Map_0 | 75 | 921.41 | (246,321) |
|  | Assembled Map_1 | 88 | 1,242.40 | (11,95) |
|  | Assembled Map_2 | 30 | 531.65 | (225,255) |
|  | Assembled Map_3 | 44 | 759.16 | (181,228) |
|  | Assembled Map_4 | 107 | 1,699.60 | (87,194) |
| Corrected Rmaps | Assembled Map_0 | 102 | 1,397.60 | (225,322) |
|  | Assembled Map_1 | 237 | 3,348 | (8,230) |
| Error-free Rmaps | Assembled Map_0 | 60 | 808.74 | (185,239) |
|  | Assembled Map_1 | 91 | 1,100.5 | (241,324) |
|  | Assembled Map_2 | 104 | 2,474.4 | (19,185) |

### Experiments with OMSim Data

To present the robustness of our method and its applicability across datasets, we conducted experiments on synthetic data from an optical map simulating software called OMSim [14]. As described in Section Datasets, we generated eight datasets of synthetic optical maps by varying the insertion and deletion error rates.

In the first experiment, we fixed the False Negative rate at 15% and varied the False Positive rate between 0.5, 1.0, 2.0 and 5.0 respectively. For each of the four datasets, we align each Rmap (using the Valouev aligner) before and after error correction to the reference optical map obtained using the same restriction enzyme and report the percent of Rmaps whose alignment S-score increased after error correction and the mean increase in the S-score. We note that for each Rmap, the aligner returns the highest scored alignment and the score represents how closely the Rmap aligns to the reference genome-wide optical map. Table 6 summarizes the results from this experiment. We observe that the eficiency of error-correction improves as the FP rate is initially increased. When the FP rate reaches a high value of 5, the eficiency of error-correction drops. The mean S-score improves by more than 9 (∼ 10%) when the FP rate is reasonable.

In the second experiment, we first fix the False Positive rate at 1.0 and vary the False Negative rate between 5%, 15% and 25%. We then fix the False Positive rate at 2 and vary the false negative rate between 5%, 15% and 25%. We report the same results as the previous experiment. Table 7 shows the results. Similar to the previous experiment, we find that the eficiency of error correction improves as the FN rate increases from 5% to 25%. The error correction improves the quality of a high percentage of Rmaps (> 70%) for all values of parameters.

### Experiments with Real Data

Table 5 summaries the results of running cOMet on the plum and goat datasets. The plum and goat datasets do not contain error-free Rmaps. Therefore, we are restricted to reporting the number of corrections made and the improvement to the *S*-score. In order to compute the *S*-score before and after error correction, we generated an *in silico* digested genome-wide optical map from the reference genome and aligned both the uncorrected and corrected Rmap to the genome-wide optical map. If it aligned to multiple positions then we considered the alignment position where the corrected Rmap aligned with greatest *S*-score, and considered the difference in the *S*-score when the uncorrected and corrected Rmap aligned to that position. However, we note that this process is error prone because of the fragmented nature of the draft genomes and possible misassemblies present in the genomes. We observed that the *S*-score after error correction improved for 78% of the plum Rmaps and 99% of the goat Rmaps. Figures 4 and 5 show the histograms of the distribution of *S*-scores, before and after error correction. For the plum genome, the mean *S*-score improved from 8.60 before error correction, to 14.72 after error correction (a 71% improvement in the score) while for the goat genome, it improved from 9.38 before correction to 16.97 after correction (a 80.92% improvement in the score).

**Table 4.** Results on the data simulated from *E*.*coli* K-12 MG 1655. The data was simulated according the algorithm described in Datasets. This simulation resulted in 2,505 Rmaps, containing 7,485 deletion and 554 insertion errors.

|  |  |
| --- | --- |
| Total no. of insertion errors corrected | 556 |
| TPR of corrected insertions | 82.49 % (457) |
| FPR of corrected insertions | 0.21 % (99) |
| Total no. of deletion errors corrected | 5,894 |
| TPR of corrected deletions | 77.38 % (5,792) |
| FPR of corrected deletions | 0.25 % (102) |

**Table 5.**
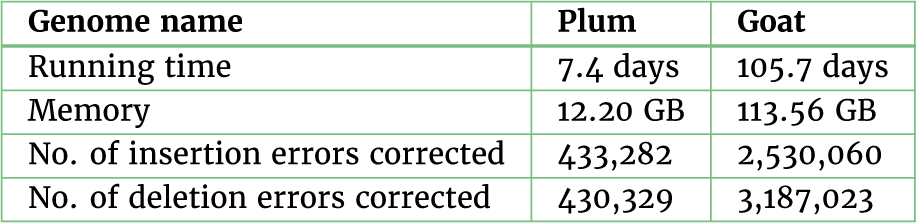
Results on the Rmap data of plum and goat genomes. Peak memory was measured as the maximum resident set size as reported by the operating system with suficient RAM to avoid paging. Running time is the user process time, also reported by the operating system.

**Table 6.**
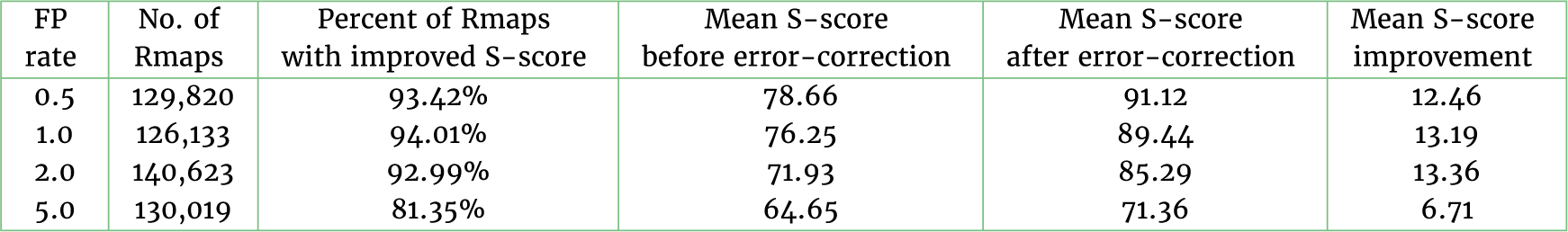
Table showing the eficiency of error correction when the False Positive rate is varied. The False Negative rate is fixed at 15%.

| FP rate | No. of Rmaps | Percent of Rmaps with improved S-score | Mean S-score before error-correction | Mean S-score after error-correction | Mean S-score improvement |
| --- | --- | --- | --- | --- | --- |
| 0.5 | 129,820 | 93.42% | 78.66 | 91.12 | 12.46 |
| 1.0 | 126,133 | 94.01% | 76.25 | 89.44 | 13.19 |
| 2.0 | 140,623 | 92.99% | 71.93 | 85.29 | 13.36 |
| 5.0 | 130,019 | 81.35% | 64.65 | 71.36 | 6.71 |

**Table 7.** Table showing the eficiency of error correction when the False Negative rate is varied.

| FP rate | FN rate | No. of Rmaps | Percent of Rmaps with improved S-score | Mean S-score before error-correction | Mean S-score after error-correction | Mean S-score improvement |
| --- | --- | --- | --- | --- | --- | --- |
| 1.0 | 5% | 142,684 | 87.36% | 81.32 | 87.62 | 6.3 |
|  | 15% | 126,133 | 94.01% | 76.25 | 89.44 | 13.19 |
|  | 25% | 123,252 | 96.09% | 70.24 | 87.44 | 17.20 |
| 2.0 | 5% | 148,912 | 89.23% | 76.54 | 84.47 | 7.93 |
|  | 15% | 140,623 | 92.99% | 71.93 | 85.29 | 13.36 |
|  | 25% | 130,763 | 93.02% | 66.95 | 81.65 | 14.7 |

**Figure 4.**
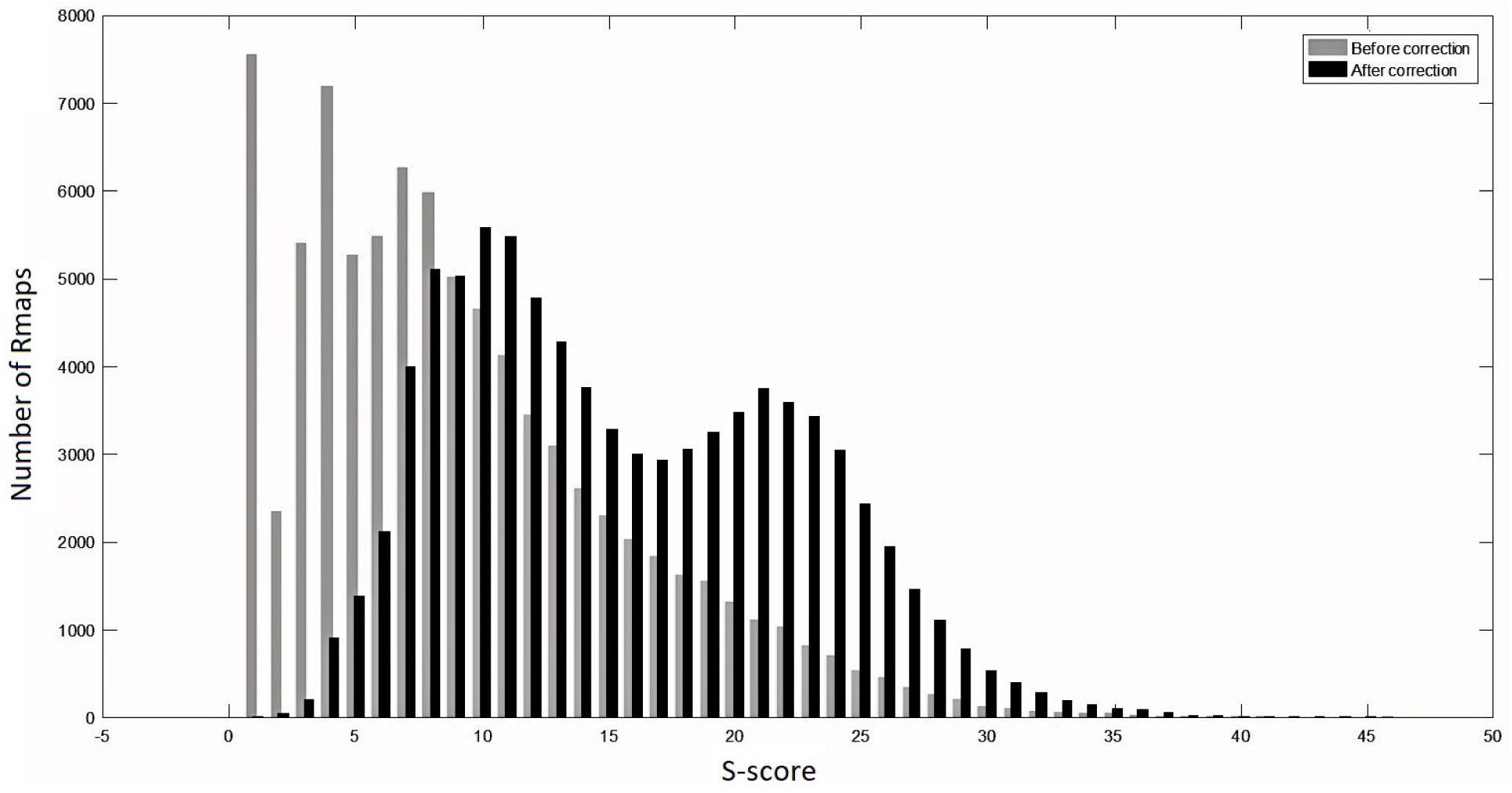
Alignment scores of Rmaps from plum genome with the reference optical-map. Before error correction, the *S*-score had a mean of 8.6 with standard deviation 6.49. After error correction, the mean *S*-score improved to 14.72, with standard deviation 6.72.

**Figure 5.**
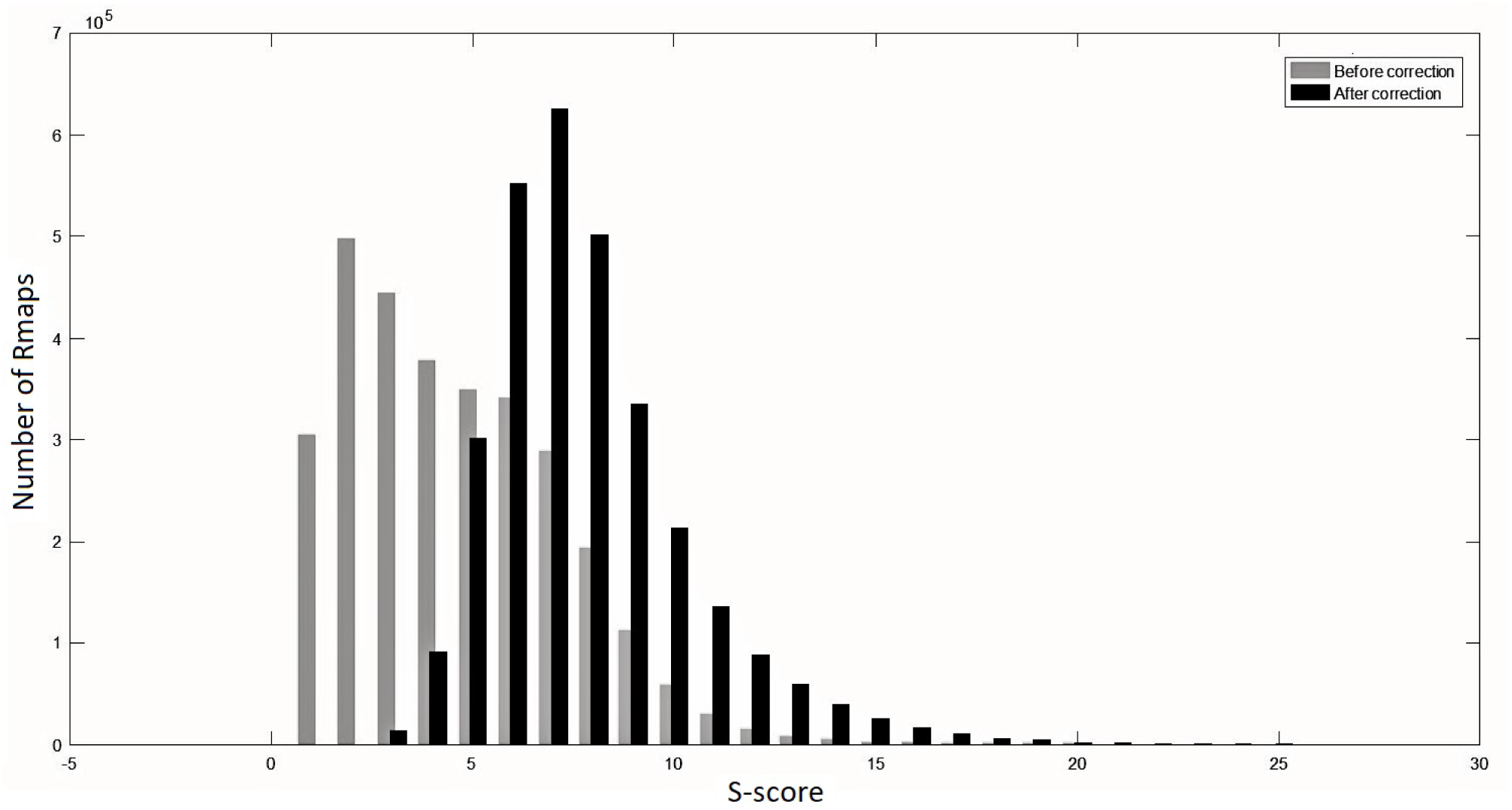
Alignment scores of rmaps from goat genome with the reference optical-map. The mean and standard deviation of the *S*-scores before error correction were 9.38 and 6.54, respectively. After error correction, the mean *S*-score improved to 16.97 with a standard deviation of 6.21.

**Figure 6.**
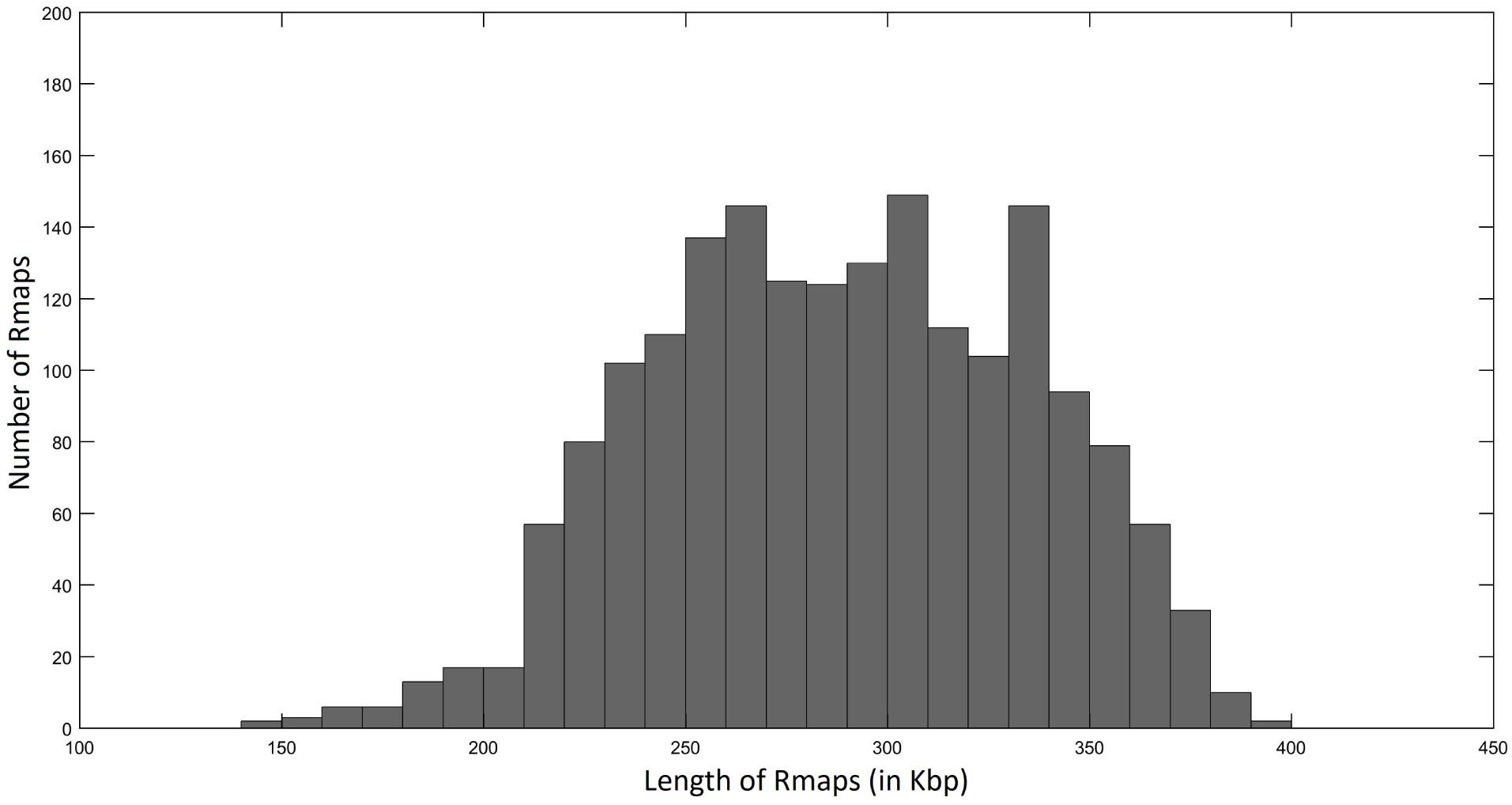
Distribution of Rmap lengths whose S-score increased after error correction. The Rmaps are simulated from the Ecoli K-12 substr. MG 1655 as explained in the text.

**Figure 7.**
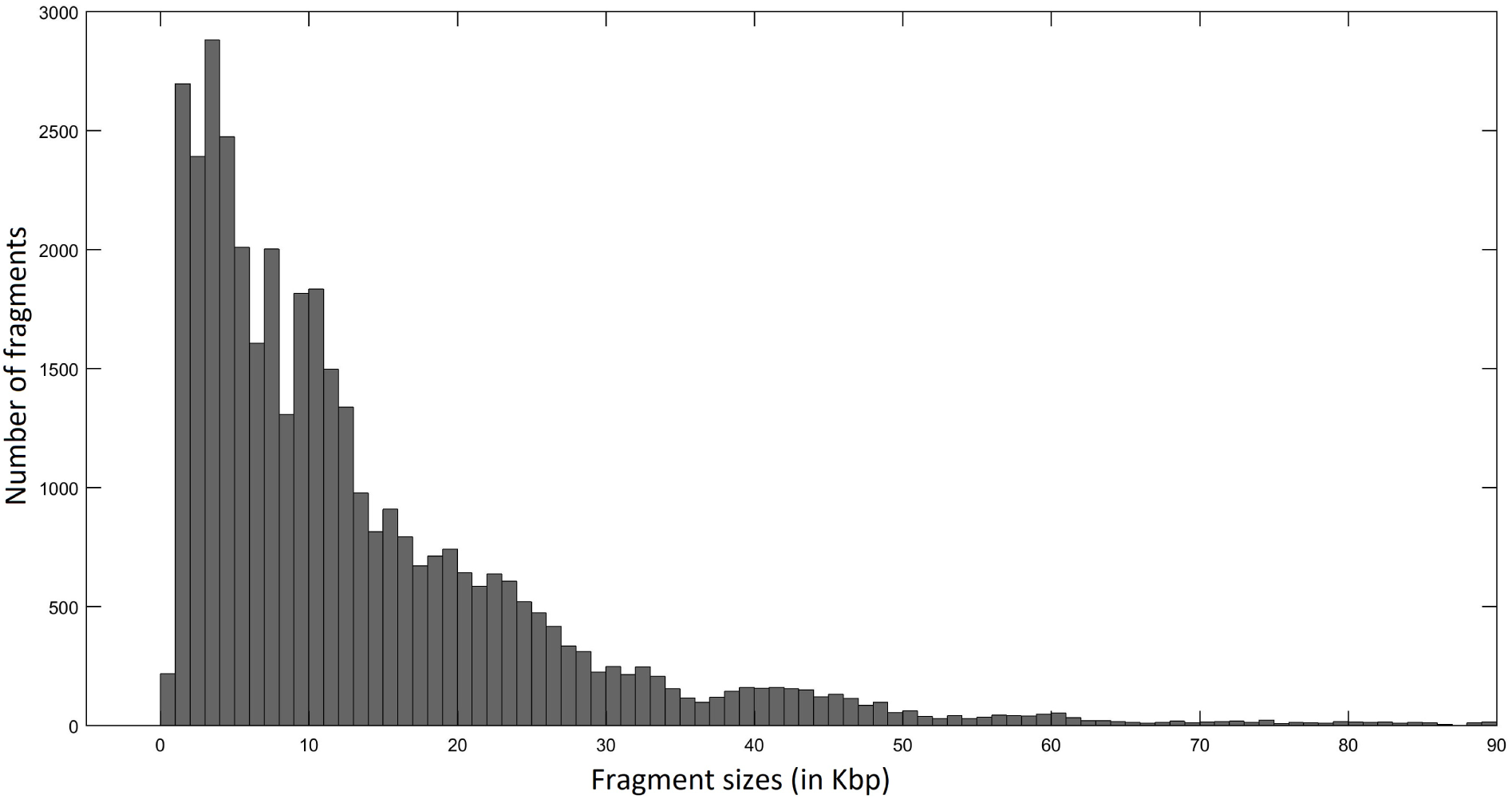
Distribution of fragment sizes of Rmaps whose S-score increased after error correction. The Rmaps are simulated from the Ecoli K-12 substr. MG 1655 as explained in the text.

We also measured the *genome coverage*, i.e. the fraction of the genome covered by at least one Rmap, for both the original Rmaps and the corrected Rmaps as follows. First we aligned all Rmaps to the genome-wide optical map and then picked the best alignment for each original Rmap and each corrected Rmap. Based on these alignments we then computed the fraction of the genome covered by at least one original Rmap and the fraction of the genome covered by at least one corrected Rmap. On the goat genome the genome coverage was 73.08% before correction and it increased to 84.56% after correction. The increase in genome coverage shows that our method is able to correct Rmaps from across the genome. Furthermore it shows that even if Rmaps could not originally be reliably aligned to some regions of the genome, our method is sensitive enough to recover similar Rmaps from these regions and thus after correction the fraction of the genome covered by aligned Rmaps is higher. For the plum, the genome coverage dropped negligibly from 99.01% before error correction to 98.85% after error correction(which is less than 1% of the genome size).

In addition, as shown in Table 5, the running time and peak memory usage was recorded for the plum and goat genome. Although these experiments have significant running times, (7.4 and 105.7 CPU days for plum and goat, respectively) these figures are not prohibitive given that this computation can easily be parallelized since the error correction process for each Rmap is independent. For example, we ran the goat genome on 20 machines and thus, it required a total of 126.84 hours for all Rmaps to be corrected. In addition, we note that error correction of a dataset will likely only be done once for any dataset so 5.2 human days for a large genome is not unreasonable. Lastly, the peak memory usage was 12.20 GB and 113.56 GB, for plum and goat, respectively, and thus, cOMet is able to run on any modern server.

Next, we ran experiments on the human dataset. Again, since we do not possess the error free Rmaps corresponding to the raw Rmaps for this dataset, we follow a similar evaluation method as in the previous experiments. We performed our evaluation on a *in silico* digested human reference genome (GenBank assembly accession: GCA_000001405.15, Genome Reference Consortium Human Build 38) using BspQI, which was the restriction enzyme that was used for generating the Rmap data. cOMet improves the S-score of 74.78% of the RMaps. The average S-score improves from 85.96 before error correction to after error correction.

## Conclusion

Error correction of high-throughput sequencing data has become an imperative pre-processing step in genome assembly since 2008 when Chaisson and Pevzner showed the dramatic improvement it can have on the quality of the assembly [3, 8, 20]. For example, after error correction the contig N50 size of an assembly of *Rhodabacter sphaeroides* improved from 233 bp to 7,793 bp using the same assembler [20]. Due to this inarguable benefit on genome assembly, many methods have been developed for error correction of sequence reads, including BFC [11], Coral [19], EULER [17, 3] and Reptile [27]. Unfortunately, even though there has been a massive effort into error correction of sequence data, there currently does not exist a publicly released method for error correction of Rmap data—a method that would likely improve the quality of genome-wide optical map assemblies, and allow such assemblies to be computed with greater eficiency.

In this paper, we presented cOMet, an error correction method for Rmap data, and demonstrate that it corrects and improves the quality of a high percentage of Rmaps in both the simulated and real datasets. As previously discussed, Rmap data is subject to high error rates. In addition to insertion and deletion errors, they contain sizing errors which necessitates the use of dynamic programming algorithm for pairwise alignment, and subsequently, assembly. By correcting a significant number of errors in Rmap data, cOMet can make it possible to use faster alignment methods [15, 10, 13], and explore the development of more eficient Rmap assembly algorithms.

## Availability of source code

The cOMet software is written in C++ and is publicly available under GNU General Public License at https://github.com/kingufl/cOMet

## Availability of Supporting Data

The optical mapping data for plum and goat is publicly available and can be accessed from their respective manuscripts. The simulated data for *E*.*coli* is provided in the github repository along with the python scripts used to generate it.

## Acknowledgements

KM, DW, MM and CB were funded by the National Science Foundation (1618814) and LS was funded by Academy of Finland (grants 284598 (CoECGR), 308030, and 314170).

## Supplementary material

In Figure 6 We show the distribution of lengths of Rmaps whose S-score increases after error correction. From the distribution we can tell that our method is able to error correct Rmaps of all sizes. We also show the distribution of fragment sizes from Rmaps whose score increases after error correction in Figure 7.

